# Pupil-size sensitivity to listening demand depends on motivational state

**DOI:** 10.1101/2023.07.27.550804

**Authors:** Frauke Kraus, Jonas Obleser, Björn Herrmann

## Abstract

Motivation plays a role when a listener needs to understand speech under acoustically demanding conditions. Previous work has demonstrated pupil-linked arousal being sensitive to both listening demands and motivational state during listening. It is less clear how motivational state affects the temporal evolution of the pupil size and its relation to subsequent behavior. We used an auditory gap-detection task (N=33) to study the joint impact of listening demand and motivational state on the pupil-size response and examine its temporal evolution. Task difficulty and a listener’s motivational state were orthogonally manipulated through changes in gap duration and monetary-reward prospect. We show that participants’ performance decreased with task difficulty, but that reward prospect enhanced performance under hard listening conditions. Pupil size increased with both increased task difficulty and higher reward prospect, and this reward-prospect effect was largest under difficult listening conditions. Moreover, pupil-size time courses differed between detected and missed gaps, suggesting that the pupil response indicates upcoming behavior. Larger pre-gap pupil size was further associated with faster response times on a trial-by-trial within-participant level. Our results reiterate the utility of pupil size as an objective and temporally sensitive measure in audiology. However, such assessments of cognitive-resource recruitment need to consider the individual’s motivational state.

**Significance statement:** The literature is inconclusive to what degree pupil size may be representing the interaction of listening task demand and an individual’s motivational state. Using an auditory gap-detection task, we manipulated both the degree of listening demand and a person’s motivational state, and investigated the temporal evolution of the pupil-size response and its relation to behavior. We find that the pupil size represents the interaction of demand and motivational state. These results highlight the importance of considering a person’s motivational state when using pupil size as a clinical measure. Pupil size appears as a key tool when assessing the impact of motivational state on the recruiting of cognitive resources.

## Introduction

Many listening situations in everyday life are characterized by speech that is masked by background sounds (e.g., music or other people talking). Communicating under such circumstances requires the allocation of cognitive resources, captured by the construct of listening effort (Eckert et al., 2016; Herrmann & Johnsrude, 2020; Peelle, 2018; Pichora-Fuller et al., 2016). Critically, the degree to which a person invests cognitive resources to achieve a goal increases with increasing situational demands as long as resources are available and the person is sufficiently motivated to achieve the goal (Brehm & Self, 1989; Richter et al., 2016). That is, a person may disengage and give up listening if the situational demands to achieve listening success exceed the person’s available resources (Brehm & Self, 1989; Herrmann & Johnsrude, 2020; Pichora-Fuller et al., 2016; Richter et al., 2016). A person may also disengage from listening, or not engage in the first place, if the reward associated with listening success is too low relative to the experienced or anticipated mental costs (Brehm & Self, 1989; Herrmann & Johnsrude, 2020; Matthen, 2016; Pichora-Fuller et al., 2016; Richter et al., 2016). Any scientifically and clinically useful measure of listening effort must therefore be sensitive to both cognitive load and motivation for it to be useful (Richter et al., 2016).

Listening effort can be measured through subjective self-reports and physiological measures (Carolan et al., 2022; Pichora-Fuller et al., 2016). Furthermore, behavioral outcomes such as changes in speech understanding, accuracy, and response time can be associated with listening effort (Houben et al., 2013; Piquado et al., 2012), but do not directly reflect listening effort (McGarrigle et al., 2014; Pichora-Fuller et al., 2016). Pupil size is often used as a physiological measure to index listening effort (Winn et al., 2018; Zekveld et al., 2018). Physiologically, a key driver of pupil size is noradrenergic neuromodulation emerging from the locus coeruleus (LC) in the brainstem (Joshi et al., 2016; Joshi & Gold, 2020). The LC in turn is sensitive to cognitive factors, such as attention (Vazey et al., 2018), and helps to optimize task performance and engagement (Aston-Jones & Cohen, 2005).

From a more audiological point of view, pupil-linked arousal is a relatively unspecific response to cognitive load during listening. For example, several speech manipulations have been shown to lead to increased pupil size: noise reduction of hearings aids being switched off (Ohlenforst et al., 2018); poor spectral resolution (Winn et al., 2016); lowered signal-to-noise ratio in speech (Koelewijn et al., 2012; Zekveld et al., 2010); syntactic complexity (Wendt et al., 2016); or semantic ambiguity (Kadem et al., 2020).

Investing cognitively under challenging listening conditions requires a listener to be motivated (Herrmann & Johnsrude, 2020; Kahneman, 1973; Pichora-Fuller et al., 2016). Motivation is often manipulated experimentally through social influences (e.g., evaluation threat: Carrillo-de-la-Peña & Cadaveira, 2000; Gilzenrat et al., 2010; Zekveld et al., 2019; perceived competence: DeWall et al., 2011; Hodgetts et al., 2019; McAuley et al., 2012) or monetary incentives (Bijleveld et al., 2009; Carolan et al., 2021; Cole et al., 2022; Gilzenrat et al., 2010; Koelewijn et al., 2018, 2021; Mirkovic et al., 2019; Zhang et al., 2019). Monetary reward prospect can improve speech perception (Bianco et al., 2021) and increase self-reported listening effort (Carolan et al., 2021). However, other work finds that monetary reward prospect does not affect pitch discrimination (Richter, 2016) or speech perception (Koelewijn et al., 2018, 2021).

A meta-analysis suggests that physiological measures of listening effort are more sensitive to monetary reward manipulations than subjective or behavioral measures (Carolan et al., 2022). Indeed, whereas some work shows that monetary reward prospect increases pupil size and performance (Van Slooten et al., 2018; Zhang et al., 2019), other work suggests that monetary reward prospect increases only physiologically measured listening effort but does not affect behavior (pupil size: Bijleveld et al., 2009; Koelewijn et al., 2018; cardiovascular reactivity: Richter, 2016). In addition, some research suggests that motivation may affect behavioral performance and physiological measures of listening effort specifically under hard, but less under easy listening conditions (Bijleveld et al., 2009; Zhang et al., 2019), consistent with motivation-effort frameworks (Brehm & Self, 1989; Herrmann & Johnsrude, 2020; Pichora-Fuller et al., 2016; Richter et al., 2016). However other research finds that motivation impacts listening effort measured through pupillometry under both easy and hard conditions (Koelewijn et al., 2018).

Critically, previous research has mainly relied on speech materials to investigate how motivation affects physiological measures of listening effort and behavior (Bijleveld et al., 2009; Koelewijn et al., 2018, 2021; Zekveld et al., 2019; Zhang et al., 2019, 2022). However, speech materials are inherently variable in content and temporal evolution which makes it harder to pinpoint how motivation, listening effort, and behavioral performance interact during listening. In the present study, we therefore use an auditory gap-in-noise detection task for which the timing of listening demand and the degree of demand can be manipulated tightly (through changes in gap timing and duration).

We seek to investigate, first, how reward prospect and listening demand jointly impact pupil-linked arousal and behavioral outcomes. Second, we will explore the temporal evolution of reward prospect on pupil size. Lastly, we aim to characterize to what extent pupil size predicts upcoming behavior.

## Material and Methods

### Participants

We recruited thirty-three younger adults (age range: 18-31 years; mean = 24.4 years; SD = 3.78 years; 7 males and 24 females; all right-handed) from the participant database of the Department of Psychology at the University of Lübeck. All of them reported no history of neural disorders nor hearing problems.

Participants gave written informed consent prior to participation and were financially compensated with 10€/hour or received course credits. In the current study, motivation was manipulated through financial rewards and participants could thus earn additional 10€ depending on their behavioral performance. The study was conducted in accordance with the Declaration of Helsinki and was approved by the local ethics committee of the University of Lübeck.

### Experimental environment

Participants were seated in a comfortable chair in a sound-attenuated booth. Participants placed their head on a chin rest positioned at about 70 cm distance from an LED computer monitor (ViewSonic TD2421, refresh rate 60 Hz). The experimental stimulation was controlled by a desktop computer (Windows 7) running Psychtoolbox-3 in MATLAB and an external RME Fireface UC sound card. Sound was delivered binaurally via in-ear headphones (EARTONE 3A, 3M). Responses were given via a four-button response box (The Black Box Toolkit, Sheffield, UK), using only one of the four buttons.

### Experimental Design

Participants listened to 5.2-s white-noise sounds that each contained one gap. The gap could occur at one of 70 randomly selected time points between 2.2 and 4.2 s after noise onset (linearly spaced; Figure 1). Participants were instructed to press a button on a response box as soon as they detected the gap (Henry et al., 2014, 2017; Henry & Obleser, 2012; Herrmann et al., 2023). Noise sounds were presented at 50 dB above a participant’s sensation level estimated using a methods of limits procedure (Herrmann et al., 2018). During the presentation of the white noise, a fixation cross was presented on the screen. Participants were asked to fixate their eyes on the fixation cross during the experiment.

**Figure 1.**
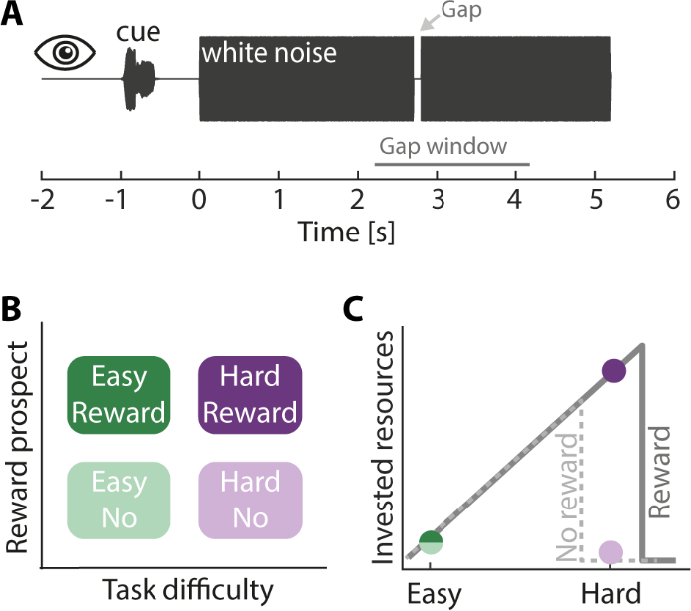
Experimental design. **A**: Auditory gap-detection task: Participants’ task was to detect a gap within a 5.2 s white-noise sound. The gap occurred at a random time between 2.2–4.2 s post noise onset (gap window, marked by the gray line). **B**: A 2 (Task difficulty) × 2 (Reward prospect) design was used. Task difficulty was manipulated by changing the duration of the gap (hard: duration was titrated to 65% detection performance; easy: twice the titrated duration). An auditory cue presented 1 s before each trial indicated whether the upcoming trial was paired with no reward or reward. **C**: Hypothesis: According to the Motivation Intensity Theory (gray lines; Brehm & Self, 1989; Richter, 2016), motivation influences cognitive resource recruitment when task difficulty is high (hard), but not when task difficulty is low (easy). In hard settings, participants should only invest cognitively when being motivated to succeed (solid line) but give up when being less motivated (dashed line). Colored dots show our hypothesis.

A 2 × 2 experimental design was used in the current study, such that we manipulated task difficulty (easy, hard) and reward prospect (no reward, reward). In detail, task difficulty was manipulated by presenting participants with white noises containing a near-threshold (hard) or a supra-threshold gap (easy). For the hard condition, gap duration was individually titrated to about 65% gap-detection performance in training blocks prior to the main experimental blocks (4–6 training blocks of 2 min each). For the easy condition, the estimated gap duration was doubled (Kraus et al., 2023).

Monetary reward prospect was manipulated such that half of the trials were paired with no reward, whereas the other half were paired with the possibility to receive an additional financial reward based on the participant’s performance on these trials. Specifically, after the experiment, three trials for each of the two difficulty levels were chosen from the reward trials (Cole et al., 2022; Teoh et al., 2020; Tusche & Hutcherson, 2018). Participants could gain 10€ in addition to their hourly compensation rate if their average performance across these six trials was above 80%.

Trials were presented in 14 blocks, each containing 20 trials. In half of the blocks, task difficulty of the 20 trials was easy, whereas in the other half of the blocks, task difficulty of the 20 trials was hard. Written information about a block’s task difficulty (easy or hard) was presented at the beginning of each block. Hence, participants had prior knowledge about whether trials contained a gap that is easy or hard to detect. Easy and hard blocks alternated, and the starting task difficulty was counterbalanced across participants.

Ten of the 20 trials per block were no-reward trials, whereas the other 10 trials were reward-prospect trials. No-reward and reward-prospect trials were presented in random order within each block. An auditory cue was presented 1 s prior to each white noise sound that indicated whether a trial was a no-reward or a reward-prospect trial. Auditory cues consisted of a guitar and a flute sound. The pairing of guitar and flute sounds to no-reward and reward-prospect trials was counterbalanced across participants. Training prior to the experiment was conducted to familiarize participants with the cue-reward association. Overall, participants listened to 70 white-noise sounds per Task difficulty (easy, hard) × Reward prospect (no reward, reward) condition, resulting in 280 trials in total.

After the experiment, participants completed a questionnaire regarding their use of the reward cues to examine whether participants were familiar with the cue-reward association during the experiment. They first indicated which auditory cue was associated with reward trials. We then asked them to rate the following statement (translated from German): “I used the auditory cues (guitar and flute) to distinguish between important and unimportant conditions” on a 6-point scale with the following labels: “strongly disagree”, “disagree”, “somewhat disagree”, “somewhat agree”, “agree”, “strongly agree”. Participants rated this statement twice, separately for the easy and the hard condition.

### Analysis of behavioral data

Any button press within 0.1 to 1 s after gap onset was defined as a hit (coded 1). Trials for which no button was pressed within this time window were considered a miss (coded 0). Response time was calculated as the time between gap onset and the button press.

Linear mixed-effect modeling using R (v4.1.2), with the packages lme4 and sjPlot, was used to analyze the influence of task difficulty and reward prospect separately on accuracy and response time. Models included effects of task difficulty (easy, hard) and reward prospect (no reward, reward) as well as the task difficulty × reward prospect interaction. Task difficulty and reward prospect were categorial predictors and were deviation coded (i.e., -0.5 [easy, no-reward] and 0.5 [hard, reward]). To account for and test for an expected hazard effect (faster responses with later gap time) we included gap time as a regressor (Herrmann et al., 2023; Niemi & Näätänen, 1981; Nobre et al., 2007). Gap time is a continuous variable and was z-transformed prior to the analysis. Response-time data were log-transformed to obtain values closer to normal distribution. We included participant-specific random intercepts to account for individual differences in overall accuracy or response time. This resulted in the statistical model expression of the following form:

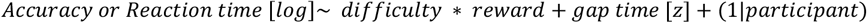

The single-trial accuracy data are binary. We thus calculated a generalized linear mixed-effect model (GLMM) with a binomial distribution and a logit link function (Tune et al., 2021). Response time (log-transformed) is a continuous variable, and we thus used a linear mixed-effect model (LMM) with a Gaussian distribution and an identity link function (Tune et al., 2021).

Our main hypothesis (derived from Motivational Intensity Theory; see Figure 1C) was that, specifically for hard trails, accuracy and response time should depend on the reward-level. Moreover, for the accuracy data, we expected performance being at ceiling for easy trials and we thus decided to elaborate the possible influence of reward in more detail for the hard condition. Hence, we employed a reduced linear mixed model using data from the two hard conditions (no reward, reward) and tested for the effect of reward prospect on accuracy and response time:

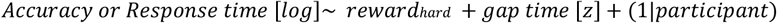

Please note that it proved not possible to include random effects for all regressors of interests due to missing convergence (Barr et al., 2013). Therefore, we decided to use only random-intercept models in order to avoid decisions about including one random effect over the other. Nevertheless, in the results section, we report if fixed effects lost their significance when including the corresponding random effect.

A paired sample t-test was used to test for effects in cue-use rating. Effect sizes for t-tests are reported as Cohen’s d (Cohen, 1988).

### Pupil data recording and preprocessing

Eye movements and pupil size of the right eye were continuously recorded using an Eyelink 1000 Plus eye tracker (SR research) at a sampling rate of 500 Hz. Data were preprocessed and analyzed using Matlab (MathWorks, Inc.). Time windows of missing data and their enclosing sharp signal increase and decrease were marked as blinks and set to NaN (‘not a number’ in Matlab; Hershman et al., 2018). If a blink occurred within 100 ms of another blink, the data points between the two blinks were also set to NaN. NaN-coded data points were linearly interpolated. Uncomplete eye lid closing can also cause artifacts in the data that do not result in missing data points (Kret & Sjak-Shie, 2019). To remove such artifacts, we calculated the median absolute deviation with a moving time window of 20 ms (Kret & Sjak-Shie, 2019). Data points that were larger than the median of the median absolute deviation + 3*IQR (inter quartile range) were marked as missing timepoints and NaN coded. If a NaN-coded time window occurred within 100 ms of another NaN-coded time window, then data points in-between were also NaN-coded. NaN-coded data points were linearly interpolated.

Pupil size can appear to change when the eyes move without an actual change in pupil size, because eye movements change the angle of the pupil relative to the eye tracking camera (van Rij et al., 2019). To account for such potential influences of eye movement on pupil size, we regressed out the linear and quadratic contributions of the x- and y-coordinates on the pupil size time course (Cui & Herrmann, 2023; Fink et al., 2021; Kraus et al., 2023). In the current study, the residual pupil size was used as the dependent measure. Pupil data were subsequently low-pass filtered at 4 Hz (Butterworth, 4th order) and divided into epochs ranging from -2 to 6.2 s time-locked to noise onset. An epoch was excluded if more than 40% of the trial had been NaN-coded prior to interpolation. If more than 50% of the trials in any condition were excluded, the full dataset of the respective participant was excluded from analysis (N=1). Pupil-size data were downsampled to 50 Hz. For each trial, the mean pupil size in the -1.6 to -1.1 s time window was subtracted from the pupil size at every time point of the epoch (baseline correction). This baseline time window was chosen to avoid contamination of the auditory cue which was presented at -1 s. For each participant, single-trial time courses were averaged, separately for each condition.

### Analysis of task difficulty and reward prospect on pupil size

To investigate the effects of task difficulty and reward prospect on pupil size, we used a similar model for the analysis as was used for the analysis of behavioral data. Non-baseline corrected pupil-size data were averaged across the time window ranging from 2.2 s (onset of gap window) to 6.2 s (end of trial) and used as dependent variable (in the literature referred to as mean pupil diameter (MPD)). Mean pupil size in the baseline time window (−1.6 to -1.1s; time-locked to noise onset) was used as a predictor to account for baseline variations in the model (Alday, 2019). Task difficulty and reward prospect were deviation-coded predictors in the model as well as their interaction. The pupil-size data were continuous and were thus z-scored:

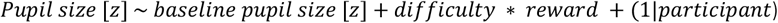

To visualize and investigate when in time the effects of task difficulty and reward prospect occurred, we calculated difference time courses. For time-resolved analyses of the main effect of task difficulty, we averaged time courses across the two reward prospect levels, separately for the easy and the hard condition. The averaged time course for the easy condition was then subtracted from the averaged time course for the hard condition. For time-resolved analyses of the main effect of reward prospect, we averaged time courses across the two task-difficulty conditions, separately for the no-reward and the reward condition. The averaged time course for the no-reward condition was then subtracted from the averaged time course for the reward condition. Every time point of the difference time courses was tested against zero using a one-sample t-test. To examine whether the effect of task difficulty differed from the effect of reward prospect, the two difference time courses (reflecting the task-difficulty and reward-prospect effects) were compared using a paired sample t-test for each time point. To account for multiple comparisons, p-values were corrected across time points using a false discovery rate of q = 5% (FDR; Benjamini & Hochberg, 1995).

For the time-resolved analysis of the interaction between task difficulty and reward prospect, we subtracted the time course for the no-reward condition from the time course for the reward condition (capturing the reward prospect effect), separately for the easy and the hard condition. Furthermore, we subtracted the time course for the easy condition from the time course of the hard condition (capturing the task-difficulty effect), separately for the no-reward and the reward condition. Every time point of the difference time courses was tested against zero using a one-sample t-test, and p-values were FDR-corrected. The reward prospect effect between easy and hard task difficulty and the task-difficulty effect between no-reward and reward were also compared for each time point (i.e., testing the interaction), again using FDR correction.

### Analysis of the relationship of pupil size and behavior

We also investigated whether a smaller pupil size increases the probability to miss the gap. To this end, we time-locked the baselined (see above) pupil-size data of every trial to the exact gap time. First, we split the trials of the hard condition into hit versus miss trials. Trials for the easy condition were not used because very few gaps were missed. To analyze whether the effect of reward prospect differs between hit and miss trials, we subtracted the time course for the no-reward condition from the time course for the reward condition, separately for hit and miss trials. Each time point of the difference time courses was tested against zero using a one-sample t-test to test at which time points pupil size was different between reward and no-reward condition. The time courses of the reward-prospect effect for the hit and miss conditions were also compared using paired sample t-tests at each time point. To analyze further whether the effect of detection (hit vs miss) differed between reward and no-reward conditions, we subtracted the time course for miss trials from the time course for hit trials, separately for reward and no-reward conditions. Again, both difference time courses were tested against zero to test at which time points hit and miss trials differed. To test whether the detection effect differed between reward and no-reward trials, the time courses of the detection effect for reward and no-reward conditions were compared with each other using paired sample t-tests. To account for multiple comparisons, p-values were corrected across time points using a false discovery rate of q = 5% (FDR; Benjamini & Hochberg, 1995).

To investigate whether pupil size prior to a gap predicts behavioral performance, we used a GLMM with the following expression:

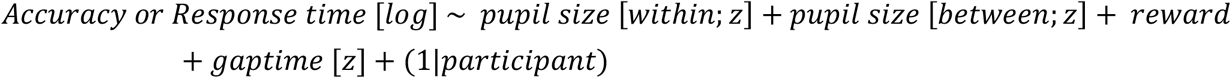

Again, only data from the hard condition were used, because of the limited number of miss trials in the easy condition. For the pupil-size regressors, pupil-size data (baselined to -1.6 to -1.1 s to noise-onset) were time-locked to the gap-onset time and averaged across the 500 ms time window prior to the gap. To disentangle associations of pupil size and behavior at the trial-by-trial state-level (i.e., within-participant) from associations at the trait-level (i.e., between-participants), we included two separate regressors associated with changes in pupil size. The between-participants regressor contained the trial-averaged pupil size per individual, whereas the within-participant regressor contained the single-trial pupil size relative to the individual mean (Bell et al., 2019; Kraus et al., 2023; Tune et al., 2021). Similar to the LMM for the behavioral analysis, we included the gap time of each trial as a regressor into the model. All continuous data (pupil-size and gap time) were z-scored across participants. Reward was deviation coded.

To further test a possible mediation effect of pupil size on the effect of reward prospect on response time, we calculated a mediation analysis in R using the package mediation. We included the following two models into the mediation analysis running 10000 iterations: (1) *Pupil size* [*z*] ∼ *reward* + *gap time* [*z*] + (1|*participant*) and (2) *Response time* [*log*]∼ *pupil size* [*z*] + *reward* + *gaptime* [*z*] + (1|*participant*). For the pupil size data, we averaged the gap-locked pupil size data across the 0.5 s before the gap.

To illustrate the association of pupil size and response time (Figure 4D), we created two groups of trials: (1) for the group of fast trials, we included all trials with response times at least +0.75 SD above the mean and (2) for the group of slow trials, we included all trials with response times at least - 0.75 SD below the mean. We chose this threshold as a good compromise between too many and too few trials per group. The grouping in fast and slow trials was done separately for reward and no-reward condition to ensure that the proportion of reward and no-reward trials does not differ between slow and fast trials. Furthermore, we only included trials such that there was no difference in gap times, relative to noise onset, between slow and fast trial responses.

### Data availability

Data and analysis scripts are available at https://osf.io/yxn96/.

## Results

### Reward prospect impacts performance under hard conditions

As expected, participants performed better and faster in the easy compared to the hard condition (Figure 2, accuracy: GLMM; odds ratio (OR) = 0.03, std. error (SE) = 0.004, p = 2.05 × 10^−153^; response time: LMM; β = 0.33, SE = 0.006, p < 2.05 × 10^−153^). An overall reward-prospect effect was present for response time but not for accuracy (accuracy: GLMM; p > 0.2; response time: LMM; β = -0.03, SE = 0.006, p = 4.89 × 10^−5^). The interaction between task difficulty and reward prospect was not significant (accuracy: OR = 1.32, p > 0.2; response time: β = -0.02, p > 0.2). The hazard effect (Niemi & Näätänen, 1981; Nobre et al., 2007) was present for response time but not accuracy (accuracy: GLMM; OR = 1.05, p > 0.2; response time: LMM; β = -0.06, SE = 0.003, p = 1.43 × 10^−66^). The later the gap appeared in a trial the faster participants responded, but the probability to detect the gap did not change as a function of gap time.

**Figure 2.**
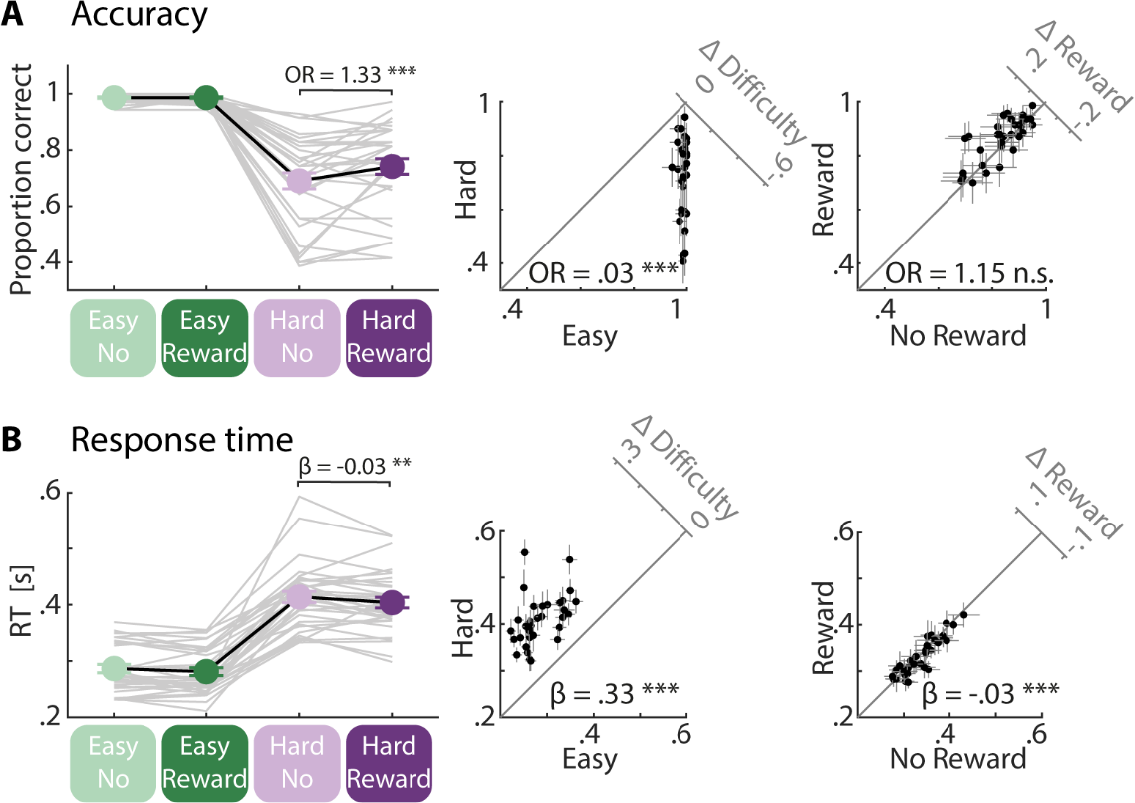
Behavioral results. **A**: Accuracy: Participants’ performance was better in the easy compared to the hard condition. In the hard condition, performance was better in the reward compared to the no-reward condition. Insets: 45-degree scatter plots showing the task-difficulty effect (left) and the reward-prospect effect (right) from linear mixed-model analysis. Difference plots (y minus x axis) are shown in upper right corners. **B**: Response time: Participants were faster in the easy compared to the hard condition and in the reward compared to the no-reward condition. In the hard condition, participants were faster in the reward compared to the no-reward condition. Insets: Same as in Panel A but for response time. Crosshairs in insets indicate 95% CI. Note that all analysis of response time used log-transformed data while figures show untransformed data for reference.

Our main interest was whether reward prospect influences behavior under the hard listening condition. Hence, we calculated a mixed-effect model-analysis using only the data from hard trials and tested for the main effect of reward prospect. We found better and faster performance in the reward compared to the no-reward condition (accuracy: GLMM; OR = 1.33, SE= 0.092, p = 7.01 × 10^−5^; response time: LMM; β = -0.03, SE = 0.011, p = 3.48 × 10^−3^). For hard trials, later gap time was associated with a faster response and a higher probability to detect the gap (accuracy: GLMM; OR = 1.08, SE = 0.037, p = 0.029; response time: LMM; β = -0.04, SE = 0.006, p = 4.89 × 10^−13^).

Interestingly, participants reported after the experiment that they used the cue indicating reward vs. no-reward trials more to adjust their listening when the task was hard compared to easy (t_32_ = 7.17, p = 3.89 × 10^−8^, d = 1.25).

### Sensitivity of pupil size to reward prospect depends on task difficulty

We first analyzed pupil-size data over a large time window from 2.2 (onset of gap time window) to 6.2 s (end of trial; time-locked to noise onset; Figure 3). Pupil size was averaged within this time window for each trial and used as dependent variable in an LMM. Pupil size was larger for the hard compared to the easy condition (effect of task difficulty: β = 0.30, SE = 0.02, p = 2.90 × 10^−83^) and for the reward compared to the no-reward condition (effect of reward prospect: β = 0.14, SE = 0.02, p = 1.06 × 10^−18^).

**Figure 3.**
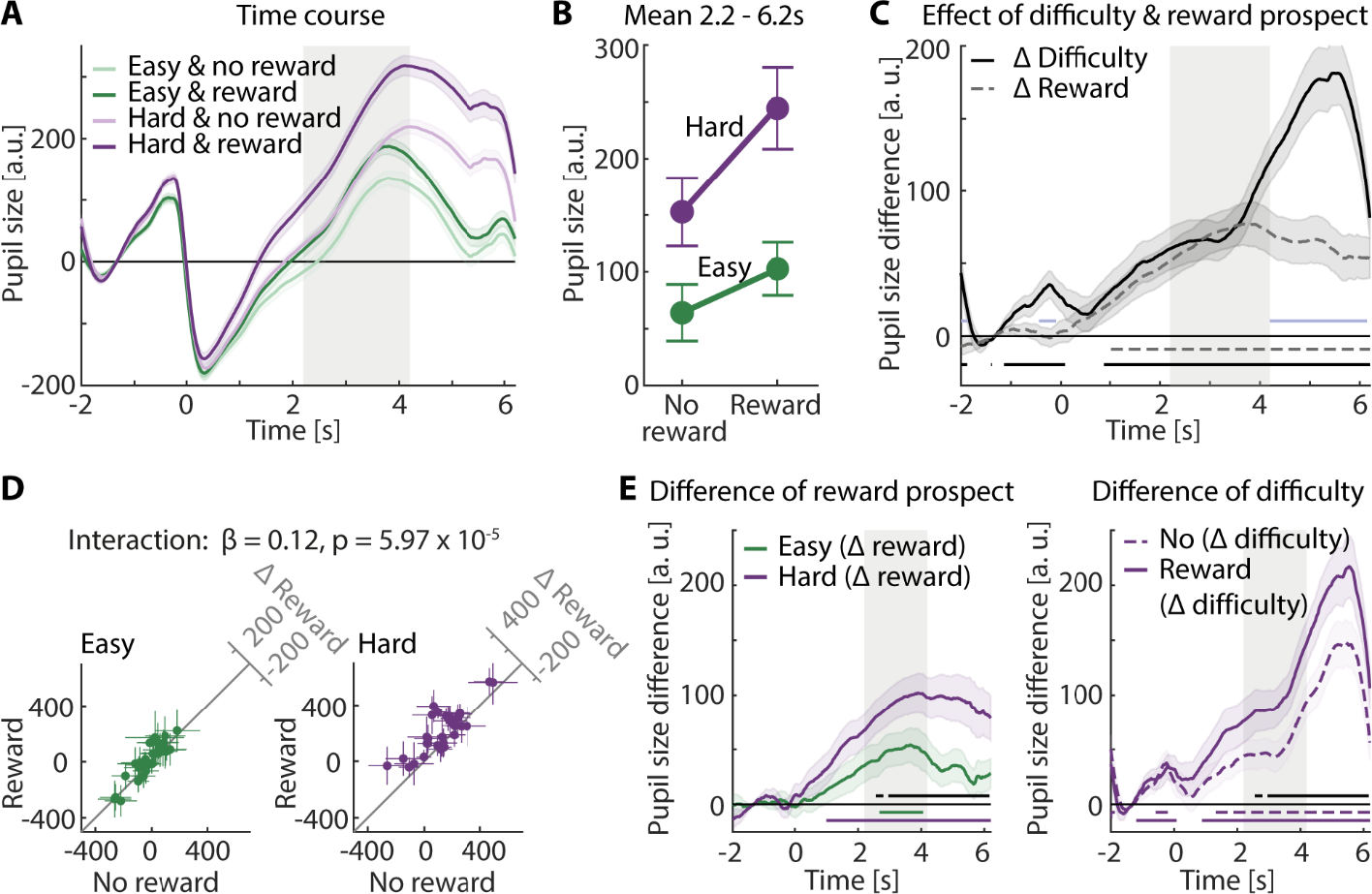
Pupil size results. **A**: Averaged pupil-size time courses across participants per condition. Error bands reflect the within-participants error. Gray areas indicate time window during which a gap could occur, from 2.2 to 4.2 s. **B**: Averaged data for 2.2–6.2 s time window. Error bars indicate the standard error of the mean. **C**: Time-resolved task-difficulty (hard minus easy) and reward-prospect (reward minus no-reward) effects. Lines at the bottom show significant time point at which main effects were significantly different from zero (black: task difficulty; gray dashed: reward prospect; FDR-thresholded) and different from each other (light blue). Error bands reflect the standard error of the mean. Gray areas indicate time window during which a gap could occur, from 2.2 to 4.2 s. **D**: 45-degree scatter plots illustrate the interaction. Left: Data from the easy condition. Right: Data from the hard condition. Colored dots show averaged pupil data per task-difficulty level, separately for each participant. The 45-degree line indicates no difference between conditions. Crosshairs indicate the 95% confidence interval (CI). Difference plots (y- minus x-axis) are shown in upper right corners. **E**: Left: Time-resolved difference of reward. Reward-prospect difference (reward minus no-reward) was calculated for each task-difficulty level and participant. Lines at the bottom show significant time points at which Δreward prospect was significantly different from zero (green: easy; purple: hard) and different from each other (black, FDR-thresholded). Right: Time-resolved difference of task difficulty. Task-difficulty difference (hard minus easy) was calculated for each reward-prospect level and participant. Lines at the bottom show significant time points at which Δtask difficulty was significantly different from zero (dashed: no-reward; solid: reward) and different from each other (black, FDR-thresholded). Error bands reflect the standard error of the mean. Gray areas indicate time window during which a gap could occur, from 2.2 to 4.2 s.

**Figure 4.**
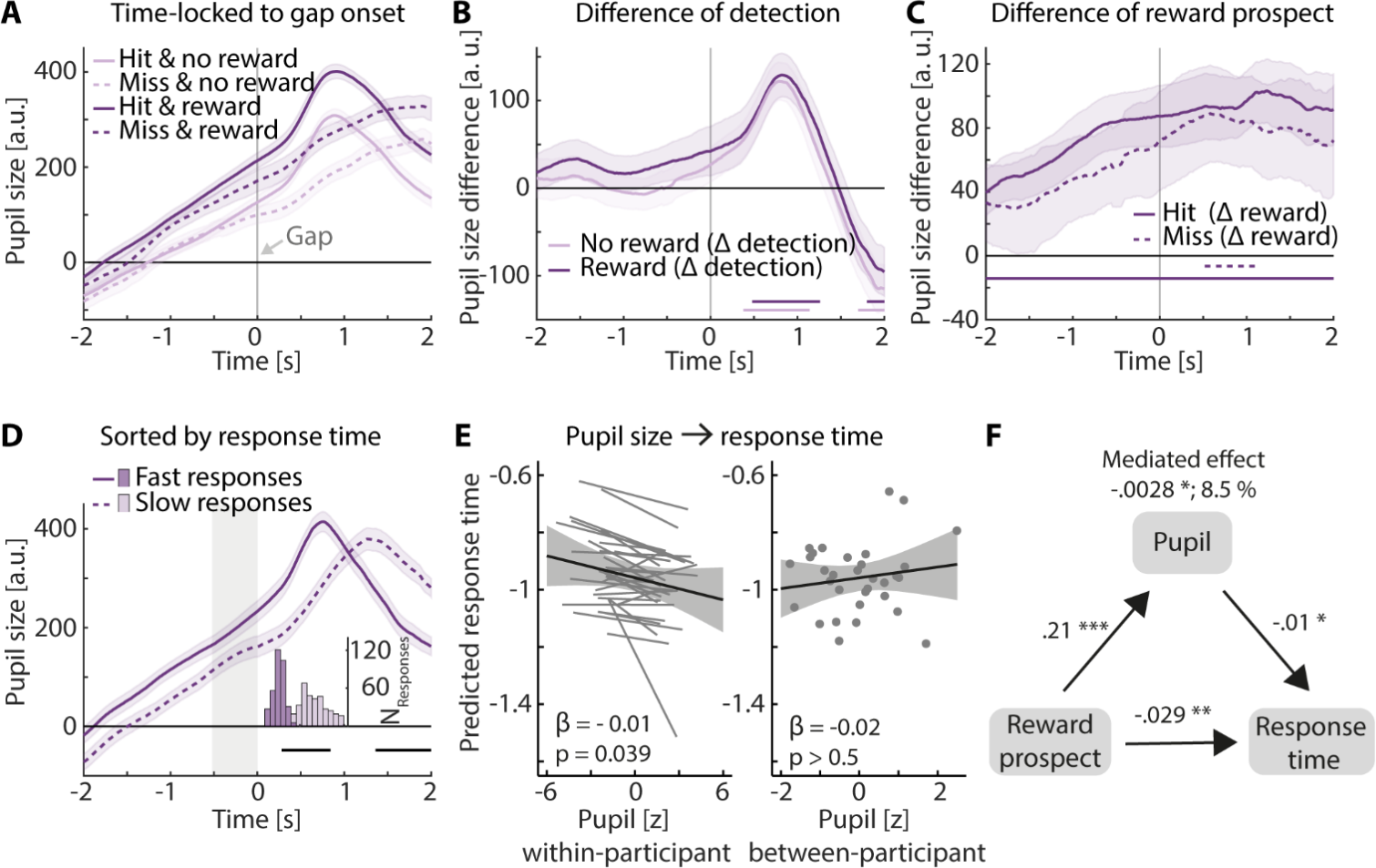
Association of pupil size and behavioral outcome. **A:** Averaged pupil-size time courses (time-locked to gap onset) for the hard condition across participants, separately for each reward-prospect condition (light vs. dark) and for hit and miss trials (solid vs. dashed lines). Error bands reflect the within-participant error. **B:** Time-resolved difference of the detection effect. Detection difference (hit minus miss) was calculated for each reward-prospect level (no-reward, reward) and participant. Lines at the bottom show significant time points at which Δdetection was significantly different from zero (light: no-reward; dark: reward). ΔDetection did not significantly differ between no-reward and reward. Error bands reflect standard error of the mean. **C**: Time-resolved difference of reward prospect. Reward-prospect difference (reward minus no-reward) was calculated for each detection level (hit, miss) and participant. Lines at the bottom show significant time points at which Δreward prospect was significantly different from zero (solid: hit; dashed: miss). ΔReward prospect did not significantly differ between hit and miss trials. Error bands reflect standard error of the mean. **D**: Average of split of pupil size data into trials with slow (dashed) and fast response times (solid). Histogram shows distribution of response times of the corresponding trials. Gray area indicates time window LMM-analysis in E. Error bands reflect the within-participant error. **E**: Effect of pupil size on response time in an LMM-analysis. Larger pupil size is associated with faster response time. Participant-specific slope for pupil-size did not improve the model but we show it here for illustrating purposes. All significant effects are based on FDR-correction. F: Mediation analysis shows that the total effect from reward on response time is partly mediated by pre-gap pupil size. * p < 0.05, ** p < 0.01, *** p < 0.001.

Critically, the Task difficulty × Reward prospect interaction was significant, indicating a larger reward-prospect effect (reward minus no-reward) when the task was hard compared to easy (LMM; β = 0.12, SE = 0.03, p = 5.97 × 10^−5^; easy reward-prospect effect: t_31_ = 2.84, p = 7.95 × 10^−3^, d = 0.502; hard reward-prospect effect: t_31_ = 5.25, p = 1.06 × 10^−5^, d = 0.927; Figure 3C).

Secondly, we calculated difference time courses to visualize and investigate in which time window the manipulation of task difficulty and reward prospect led to changes in the pupil size response. For both task-difficulty levels, pupil size was greater in the reward compared to the no-reward condition over a wide time window (easy: 2.68–4.06 s; hard: 1–6.2 s; Figure 3E left). For both reward-prospect levels, pupil size was greater in the hard compared to the easy condition over a wide time window (no-reward: -0.72– -0.06 s & 1.32–6.2 s; reward: -1.2– 0.08 s & 0.88–6.2 s; Figure 3E right). Critically, and in line with the interaction results from the temporally averaged analysis, the difference time courses (reward minus no-reward) differed across a wide time window with the task difficulty. The reward-prospect effect was larger in the hard compared to the easy condition (2.56–2.8 s and 2.96–6.16 s; Figure 3E left & right).

Furthermore, for the analysis of the time-resolved main effects, both – the task-difficulty and the reward-prospect effect – were significant almost the entire time the white noise was presented (difficulty: 0.88–6.2 s; reward: 1.02–6.2 s; Figure 3C). In addition, pupil size was larger for the hard compared to the easy condition in response to the cue, prior to the noise sound started (−1.14–0.08 s). Finally, the task-difficulty and the reward-prospect effects differed in the early cue-time window (−0.44–0.1 s), but also towards the end of the white-noise sound (4.2–6.14 s; Figure 3F).

### Larger pre-gap pupil size is associated with faster response time

We investigated whether a smaller pupil size prior to the gap is associated with a higher likelihood of missing a gap. To this end, we time-locked the pupil-size data of each trial to the respective gap time. Splitting the data in hit and miss trials enabled us to analyze whether the pupil-size time course differs depending on gap detection (Figure 4A). Pupil size was indeed larger for hit trials compared to miss trials, but only around a second after gap onset, for both reward-prospect levels (no-reward: 0.38–1.14 s; reward: 0.48–1.26 s). Towards the end of the noise sound, pupil size was larger for miss compared to hit trials for both reward-prospect conditions (no-reward: 1.7–2 s; reward: 1.8–2 s; Figure 4B). This indicates that pupil size decreased after participants detected the gap (hit), whereas, for miss trials, the pupil continued to dilate continuously until the offset of the white noise. This behavior-related effect occurred independent from reward prospect (interaction of detection and reward prospect was not significant; Figure 4B).

Likewise, the reward-prospect effect (reward minus no-reward) was independent of whether a gap was detected or not (hit vs. miss). In both cases, pupil size was larger for the reward compared to the no-reward condition. The reward-prospect effect started earlier for hit than for miss trials (hit: -2– 2 s; miss: 0.52–1.1 s; interaction of detection and reward prospect was not significant; Figure 4C).

Lastly, we examined whether the differences in pupil size between hit and miss trials may also translate to successful predictions of behavior on a trial-by-trial level. We calculated LMM models to predict accuracy or log response time from the pupil size averaged across the 0.5-s time window pre-gap onset (Figure 4D). There was no significant trial-by-trial relationship between pre-gap pupil size and accuracy for the within-participant effect (GLMM; OR = 1.02, p > 0.5) or for the between-participant effect (OR = 1.13, p > 0.4). However, within participants, trials with larger pre-gap pupil size preceded shorter response times in these trials (LMM; β = -0.013, SE = 0.006, p = 0.039; no effect at the between-participants level; β = -0.015, p > 0.5) (Figure 4E).

Since we found an effect of reward prospect on response time and pupil size, and that the pupil size and response time are associated, we asked if the effect of reward prospect on response time may be mediated by pupil size. Therefore, we ran a mediation analysis resulting in a small mediation effect of pupil size (β = -0.0028, p = 0.035, proportion mediated: 8.5 %).

## Discussion

In the current study, we investigated the extent to which task difficulty and a listener’s motivational state, here manipulated as reward prospect, influence pupil size and ensuing behavior. An auditory gap-detection task allowed us to manipulate both the task difficulty (i.e. gap duration) and precise timing of a target (gap).

Reward prospect improved response time and accuracy under difficult listening conditions. Critically, pupil dilation was greater under higher compared to lower task demands and especially so when reward prospect was high. This result emphasizes the fact that pupil size cannot be taken as an invariant read-out of listening demand but is also sensitive to an individual’s motivational state, especially under more challenging listening conditions.

### Reward prospect improves behavioral performance

Behavioral performance decreased with increasing cognitive demand, as expected based on our manipulation (see also Herrmann et al., 2023; Kraus et al., 2023). Critically, reward prospect was associated with better gap-detection accuracy and response time when listening was hard (response time was also faster under easy listening conditions; Figure 2B). Some previous work also observed reward-related performance increases in auditory tasks (Bianco et al., 2021; Mirkovic et al., 2019; Zhang et al., 2019), whereas other studies did not (Carolan et al., 2021; Koelewijn et al., 2018). Why behavioral benefits related to manipulations of motivation are observed only sometimes could be due to different tasks (mainly speech tasks compared to present study: Bianco et al., 2021; Carolan et al., 2021; Mirkovic et al., 2019; Zhang et al., 2019), different manipulation of task difficulty (speech vocoding: Carolan et al., 2021, speech rate: Zhang et al., 2019, directional microphone technique in hearing aids: Mirkovic et al., 2019, target saliency: Koelewijn et al., 2018, 2021), and/or the amount of monetary reward prospect. It appears that the maximum achievable reward has been lower in studies that did not observe reward-performance benefits (£2.50 Carolan et al., 2021; 5€ Koelewijn et al., 2018) compared to those observing such benefits (10€ in the present study; 20€ in Mirkovic et al., 2019; 50€ in Zhang et al., 2019), possibly adding to the inconstancies across studies.

We observed no statistically significant interaction between task difficulty and reward prospect on behavioral outcomes (accuracy; response speed; Figure 2). This is not easily reconciled with Motivation Intensity Theory (Brehm & Self, 1989; Richter, 2016) and also empirical listening-effort work (Mirkovic et al., 2019; Zhang et al., 2019), which suggest that motivation should exhibit the strongest benefit under harder task conditions, and possibly no effect under easy conditions. Under hard conditions, people in a low-motivation state may give up because the effort required is not worth the reward (Figure 1B), whereas under easy conditions, participants should have no problem to perform the task regardless of motivational incentives.

The observation that the reward effect was not significantly stronger under hard compared to easy listening condition, as suggested by the main effect and absence of an interaction, may be due to several reasons. During the “hard” condition under no-reward, participants had nothing else interesting to do and may have invested cognitively. The “hard” condition may also not have been hard enough for people to give up under the no-reward condition on most trials (∼69%; Figure 1C). Another reason could be that the “easy” condition still required individuals to invest effort despite ceiling performance accuracy (cf. work showing same behavioral performance but different amount of listening effort: Bentler et al., 2008; Gagné et al., 2017; Picou & Ricketts, 2014). People may thus have invested more cognitively during reward than no-reward trials even when task difficulty was easy, leading to response time benefits from reward for both the easy and the hard listening conditions.

Interestingly, although there was no significant interaction between task difficulty and reward prospect, participants’ self-reports about cue-use showed that participants indicated to use the auditory cue stronger in the hard compared to the easy condition to adjust their listening to the task. This is similar to Carolan et al (2021), who found reward effects on self-reported listening effort but not on performance.

### Pupil-size sensitivity to listening demand depends on motivational state

We show increased pupil size for high compared to low task demands, consistent with previous work (Kadem et al., 2020; Kahneman & Beatty, 1966; Koelewijn et al., 2012; Kraus et al., 2023; Ohlenforst et al., 2018; Wendt et al., 2016; Winn et al., 2016; Zekveld et al., 2010; Zekveld & Kramer, 2014; Zhao et al., 2019). We also observed that monetary reward prospect for good task performance increased pupil size (Figure 3), indicating that motivational state plays a key role in modulating pupil size. A recent meta-analysis also suggests that pupil size is sensitive to reward manipulations (Carolan et al., 2022; see also Bijleveld et al., 2009; Koelewijn et al., 2018; Manohar et al., 2017; Zhang et al., 2019), although not all studies show motivation effects on pupil size (Gilzenrat et al., 2010; Koelewijn et al., 2021).

Critically, task difficulty and reward prospect interacted: The increase in pupil size due to reward prospect was greater under hard compared to easy listening conditions (Figure 3), suggesting that the investment of cognitive resources is greater when a listener is highly motivated, especially in hard listening situations.

The interaction between task difficulty and reward prospect on pupil size is in line with Motivation Intensity Theory (Brehm & Self, 1989; Richter, 2016), in that, a higher influence of motivation on effort (pupil size) was expected in highly demanding compared to less demanding situations. Motivation Intensity Theory predicts that a person only invests cognitively when they have sufficient cognitive resources available and are motivated (Brehm & Self, 1989; Richter, 2016), otherwise they would give up listening.

Under hard conditions, people may have given up investing effort (Ohlenforst et al., 2017; Wendt et al., 2018; Zekveld & Kramer, 2014) more frequently compared to easy conditions. Under easy listening conditions, participants had sufficient cognitive resources to perform the task and were motivated to do so as indicated by the ceiling performance (Figure 2). However, our data unexpectedly show also for the easy condition that pupil size – and thus listening effort – is larger during reward compared to no-reward trials, whereas no difference was predicted by the Motivation Intensity Theory (Brehm & Self, 1989; Richter, 2016; Figure 1). It is possible that participants occasionally gave up listening under the easy, no-reward condition because they were bored or underchallenged (see Herrmann & Johnsrude, 2020; Westgate & Wilson, 2018), but then still detected the gap because it was perceptually sufficiently salient in the easy condition (consistent also with longer response times for the no reward than the reward condition). Yet, if this low-effort strategy was employed by listeners, why they would not have used it under the reward condition – where pupil size indicates higher effort – is not clear. Future research may need to further address the impact of motivation under easy listening conditions.

Not least the present results hold an important lesson for the increasingly common use of pupil dilation as a clinical measure. Pupil dilation cannot serve as a direct readout of purely physical listening demand. It must be taken into account that the internal motivational state of a person is not only influencing pupil dilation directly. Motivational state also gates or constrains the size of the effects that listening demand will exert on pupil size. Furthermore, a person’s motivational state can affect their pupil size even when influences on behavioral performance do not emerge.

### Pupil size predicts response time

The time-resolved analysis of pupil size revealed that pupil size differed depending on whether a listener detected the gap or not, whereas pupil size did not predict accuracy on a trial-by-trial level. However, we found that a larger pupil size was associated with faster response times. These results are in line with work suggesting that response time may be a more sensitive behavioral measure compared to accuracy (Houben et al., 2013; Kang et al., 2017; Weis et al., 2013) and that response time is associated with pupil-linked arousal (Schriver et al., 2018; Van Kempen et al., 2019; Wainstein et al., 2017). Using our auditory-gap detection task allowed us to investigate when in time effort is invested, revealing that pupil size predicts response time measures. This can be interpreted as higher cognitive resource investment measured by pupil size helps optimizing behavioral outcome. Time-resolved analyses may be more challenging for complex speech stimuli, for which it is less clear when effort is invested (Bijleveld et al., 2009; Gilzenrat et al., 2010; Koelewijn et al., 2018, 2021; Zhang et al., 2019). Furthermore, the present work highlights the use of single-trial models to analyze pupil-behavior-associations (Kraus et al., 2023; Tune et al., 2021).

Previous work has suggested a link between LC-NE activity and performance increases (Aston-Jones et al., 1994; Aston-Jones & Cohen, 2005; Usher et al., 1999), which may be reflected in the pupil size. Our mediation analysis revealed that a small part of the influence of reward on response time is mediated via pupil size. Hence, only a small portion of the behavioral benefit under reward can be explained via enhanced LC-NE activity that is indexed by pupil-size changes. Our data suggest that most of the reward effect directly impacts behavioral performance or via pathways different from the LC-NE pathway.

## Conclusion

The motivational state of a person influences the extent to which they invest cognitive resources under hard demands. We used an auditory gap-detection paradigm that tightly controlled task demands and target timing. Pupil size reflected stronger influences of a person’s motivational state under hard than easy listening conditions. This highlights the importance for future work to not see pupil size as a simple read-out of task demand. Moreover, pupil size is thus a key indicator when considering an individual’s motivational state in measuring cognitive investment.

## Acknowledgments

We thank Aigerim Tuichieva for assisting with the data collection. This work was supported by Deutsche Forschungsgesellschaft (DFG; grant number HE 7857/1-1) awarded to BH. BH was supported by the Natural Sciences and Engineering Research Council of Canada (Discovery Grant: RGPIN-2021-02602) and the Canada Research Chair program (CRC-2019-00156, 232733).

